# Geothermal gradients restructure metabolism in hot spring microbial communities

**DOI:** 10.64898/2026.09.13.751272

**Authors:** Danli Luo, Christaline George, Patrick K.H. Lee, Stephen B. Pointing

## Abstract

Hot springs are employed as model systems for understanding how environmental stress structures microbial communities, yet the mechanisms by which geothermal stress reorganizes community metabolism remain poorly understood. Here we combined genome-resolved metagenomics, metatranscriptomics, aqueous geochemistry, and community metabolic modelling to investigate photosynthetic biofilms across the geothermal gradient of a hot spring in Singapore. We reconstructed community taxonomic and functional composition via metagenome-assembled genomes and identified three transcriptionally coordinated ecological guilds defined by convergent metabolic strategies rather than phylogeny. Community metabolic models revealed that increasing geothermal stress reorganized nutrient exchange networks, progressively reducing metabolic redundancy and increasing dependence on reciprocal interactions among thermophilic guilds. Increasing geothermal stress redirected relative modelled elemental demand from carbon-associated metabolism towards nitrogen and sulfur metabolism, highlighting nitrogen limitation and sulfur-linked bioenergetics as defining features of stress adaptation. Electron allocation modelling further demonstrated a systems-level transition from growth-oriented biosynthesis toward catabolic energy conservation, respiratory metabolism, and oxidative stress management under geothermal stress. These coordinated metabolic shifts indicate that hot spring communities become increasingly specialized and energetically constrained near their upper geothermal limits. Overall, the findings demonstrate how environmental stress restructures microbial ecosystems through guild-level metabolic partitioning and cooperative resource exchange and establishes an integrative systems biology framework for predicting microbial ecosystem function in extreme environments.

## Introduction

Hot springs are geothermal ecosystems characterized by extreme physicochemical conditions that create challenging habitats for microbial life^1^. They have long served as model systems in microbial ecology because they occur across broad geographic scales, possess readily identified major abiotic variables, and are subject to minimal influence from metazoans^2–6^. One of the most visible forms of colonization are photosynthetic biofilms that proliferate at temperatures ≤ 75 °C^7^. However, their microbial complexity poses substantial challenges for resolving functionality in their extreme habitat. A number of photosynthetic hot spring taxa have been cultivated and aspects of their metabolism interrogated^8–10^, but the inability to replicate biofilm complexity and habitat conditions under laboratory conditions have created a major knowledge gap regarding *in situ* functionality of microbial communities^11^.

Traditional sequencing-based approaches have partly addressed cultivation challenges. Advances in metagenomics have enabled the characterization of community structure and functional potential across diverse hydrothermal environments^12–15^, as well as predictions of putative inter-dependencies based on co-occurrence and complementarity networks^16^. Metatranscriptomics is also beginning to provide greater understanding of putatively active microbial taxa for specific habitats, such as acidic geothermal sediments at Tengchong in China^17^, and phototrophic biofilms at Yellowstone National Park in the USA^18,19^. While informative, these studies have mainly performed co-occurrence analyses to qualitatively explore biotic and abiotic interactions. When multiple environmental limitations elicit overlapping metabolic responses, such analyses cannot provide clear quantitative insights on community scale interactions^20,21^. However, such multifaceted interactions are expected to be common in natural ecosystems where multiple limiting factors in addition to the primary thermal stress in hot springs, such as carbon^13,22^, nitrogen^22,23^, phosphorus^14^, trace elements^24,25^, energy sources^22,26^, and oxidants^13,27^, may act simultaneously.

Community metabolic models (CMMs) offer a potentially transformative systems biology analytical framework to more fully reveal the putative functional dynamics and interactions of microbial communities^20,28,29^. These models integrate multiple data sources that describe the entire metabolic network based on genomes, stoichiometric matrices that describe all metabolic reactions based on gene expression and nutrient availability, and gap-filling algorithms that complete missing pathways. These are used to create a mathematical representation of the putative metabolic networks for multiple organisms, allowing researchers to simulate growth and predict metabolic interactions within microbial communities^29,30^. A challenge in the adoption of CMMs lies in generating validated datasets for diverse microbial communities and effectively translating multi-omics and abiotic data into reliable model constraints^28–30^, while accounting for uncertainty in transcriptome-constrained metabolic states. Recent applications have revealed the ability of CMMs to generate major novel insight on microbial community functionality in human^29,31^, marine^20^, wastewater^32^, and food fermentation^28^ systems. We hypothesized that developing a well-constrained CMM for hot spring biofilms would reveal valuable novel insight on how hot spring communities, which are often identified as a model system for microbial ecology studies^4,16^, are functionally organized and respond to stress via the model-driven integration of metagenomes, metatranscriptomes, and aqueous geochemistry.

Here, we describe a novel systems biology approach to interrogate biofilms along a well-defined thermal and physicochemical geothermal gradient at the Sembawang Hot Spring in Singapore. We developed a CMM to integrate data from metagenome-assembled genomes with metatranscriptomics and abiotic parameters (Fig. 1). Our data-driven simulations were constrained by active microbial community composition, metabolic reaction flux constraints, and environmental constraints. The context-constrained models were extensively validated to produce robust ecological inference on predicted nutrient and electron allocation in biofilms. This allowed us to quantitatively express three novel metrics of microbial metabolism within the biofilm community: The Association Coefficient (AC) for co-consumption and cross feeding of nutrients, the Elemental Allocation Index (EAI) for defining nutrient limitations, and the Anabolic Reallocation Bias (ARB) that describes electron allocation prioritization of anabolic, catabolic, and stress response sinks. Overall, the findings illustrate how metabolic interactions among cooperating guilds facilitate metabolic re-prioritization under environmental stress.

**Figure 1.**
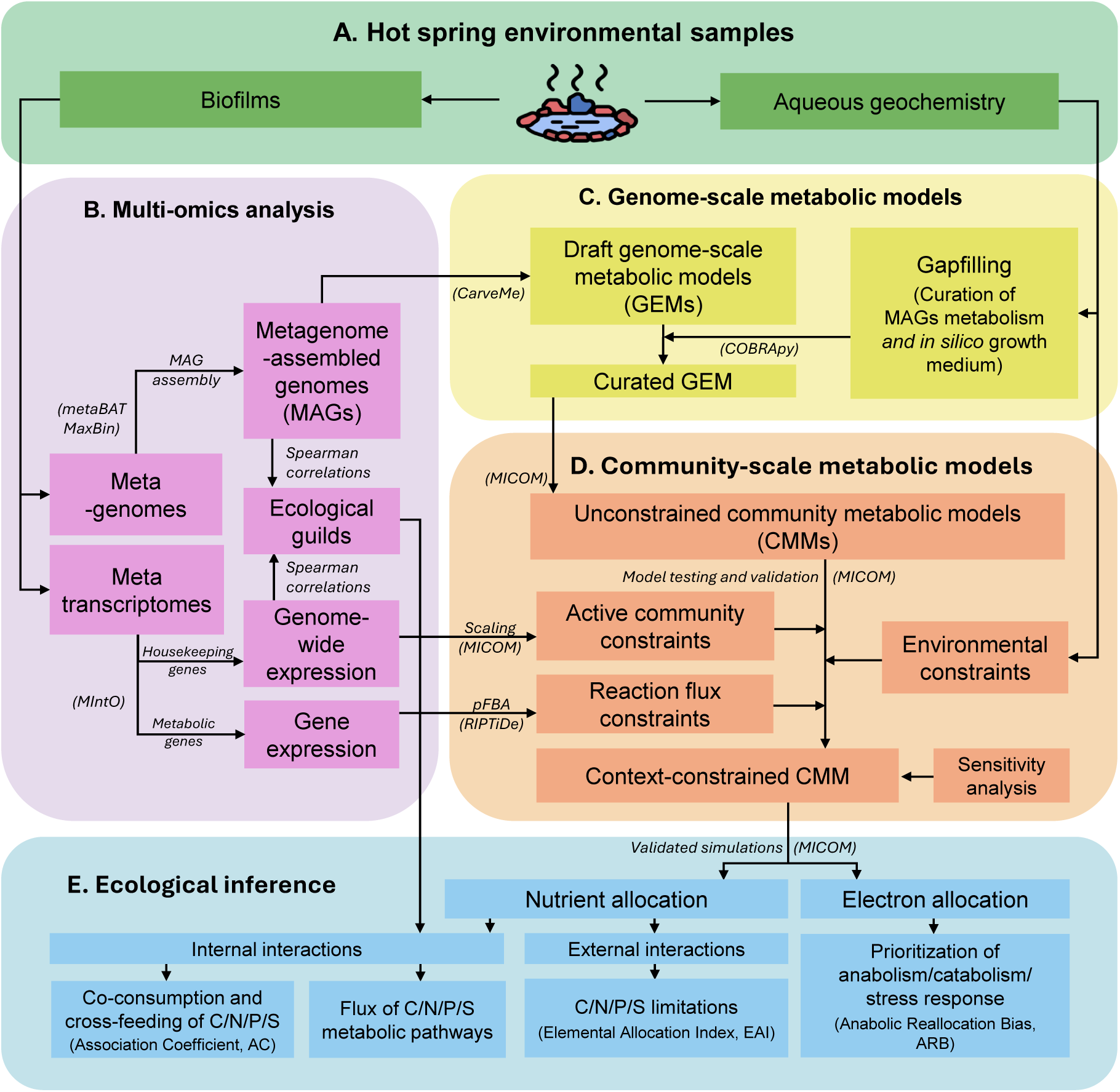
Multi-omics workflow for context-constrained metabolic modelling of hot spring biofilm communities. **A)** Environmental biofilm samples and aqueous geochemical data were collected from hot spring ecosystems. **B)** Sequencing and ecological grouping integrated with metagenomic and metatranscriptomic analyses to reconstruct community metabolism across the thermal gradient. **C)** Metagenome-assembled genomes (MAGs) were used to generate draft genome-scale metabolic models (GEMs), followed by manual curation and gapfilling under environmentally relevant growth conditions. **D)** Community-scale metabolic models (CMMs) were subsequently constructed in a MICOM framework and constrained using environmental chemistry, active community composition, and transcriptionally informed reaction bounds to generate context-constrained simulations of microbial metabolism. **E)** Validated models were then used to infer estimates of nutrient allocation and electron allocation dynamics.

## Results

### Genome-resolved reconstruction of hot spring biofilm communities

We reconstructed metagenome-assembled genomes (MAGs) spanning 20 bacterial phyla (Supplementary Dataset 1), from phototrophic biofilms sampled across a 46–61 °C geothermal gradient comprising multiple measured physico-chemical stressors (with samples referred to by their temperature for ease of identification) at the circumneutral Sembawang Hot Springs Park in Singapore (1.4343° N, 103.8225° E)^33^. Read recruitment analysis showed that 140 high- and medium-quality MAGs captured most community diversity, accounting for an average of 77% of quality-controlled metagenomic reads (range 61–88%, Supplementary Dataset 1). This refined MAGs dataset was used in all downstream analyses. Chloroflexota and Cyanobacteria dominated biofilms across the gradient, although clear geothermal preferences were evident among individual MAGs and phylogenetic groups. Other moderately abundant taxa included members of the Bacteroidota, Planctomycetota, and Pseudomonadota. Integration of these genomes with metatranscriptomes and aqueous geochemistry measurements (Supplementary Dataset 2) provided the basis for reconstruction of context-specific community metabolic models (CMMs) to investigate metabolic interactions, nutrient allocation, and bioenergetic allocation across the geothermal gradient.

### Ecological guilds partition metabolism across the geothermal gradient

Genome-wide gene expression profiling revealed strong ecological partitioning among MAGs across the gradient (Fig. 2A). Phototrophic Cyanobacteria and Chloroflexota dominated the most metabolically active fraction of the community, although distinct lineages displayed contrasting geothermal optima. Filamentous diazotrophic Cyanobacteria, including Ellainellaceae and *Fischerella*, exhibited greatest activity at moderate geothermal stress, whereas unicellular *Thermosynechococcus* and anoxygenic phototrophic Chloroflexota increased in activity toward the upper thermal range. In contrast, many heterotrophic Planctomycetota and Pseudomonadota declined in activity above 55 °C, while Desulfobacterota, Elusimicrobiota, and Nitrospirota were largely active at the most elevated geothermal stress.

**Figure 2.**
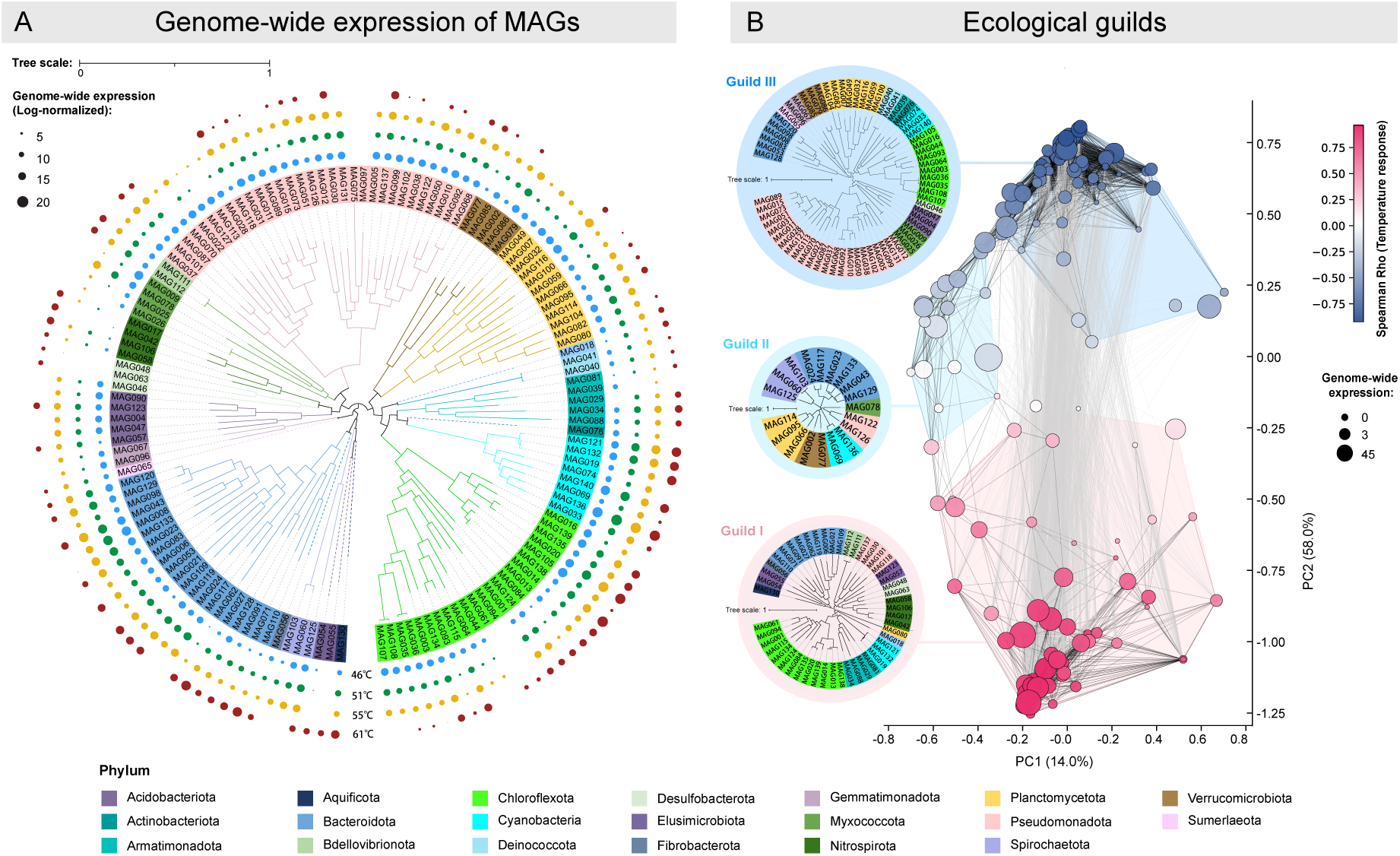
Genome-resolved transcriptional activity and ecological organisation of hot spring biofilm communities. **A)** Phylogenomic distribution and genome-wide gene expression (GE) of metagenome-assembled genomes (MAGs) recovered from hot spring biofilms. The maximum-likelihood phylogenomic tree is annotated by phylum-level taxonomy and concentric rings representing log-normalised GE derived from metatranscriptomic mapping. Point size indicates relative transcriptional activity for each MAG across the sampled geothermal gradient. Major bacterial phyla included Acidobacteriota, Chloroflexota, Cyanobacteria, Aquificota, Bacteroidota and Proteobacteria-related lineages, revealing substantial variation in transcriptional allocation among phylogenetic groups. **B)** Ecological grouping of MAGs based on genome-wide transcriptional responses to temperature stress. Principal coordinate analysis (PCoA) revealed three major ecological guilds (GI–GIII), each characterised by distinct coordinated expression profiles and interaction structures. Node size represents GE magnitude, while node colour indicates Spearman’s correlation with temperature response. Edges represent significant co-expression associations between MAGs. Insets show the phylogenetic composition of each ecological guild, illustrating that transcriptionally coherent ecological responses emerged across multiple phylogenetic lineages rather than within single taxonomic groups alone.

To examine how transcriptional organization varied across the geothermal gradient, we inferred ecological guild structure using pairwise genome-wide gene expression correlations and maximum-modularity partition analysis, while integrating Spearman correlation patterns to visualize organization across the network structure (Fig. 2B). This was evaluated and determined as superior to an abundance weighted correlation (Supplementary Dataset 2). We identified three transcriptionally coordinated guilds (GI–GIII) that reflected convergent metabolic strategies rather than phylogenetic affiliation (Fig. 2B, Supplementary Dataset 2). GI comprised thermophiles adapted to the highest geothermal stress, enriched in Chloroflexota, unicellular Cyanobacteria, and Nitrospirota which were associated with reductive sulfur cycling, hydrogen metabolism, thermophilic carbon fixation, and anaerobic organic matter turnover. GII represented a transitional nutrient-recycling guild dominated by heterotrophic Bacteroidota, filamentous Cyanobacteria, and Planctomycetota involved in polymer degradation and intermediary nutrient exchange. GIII formed an oxidative guild with a preference for lower geothermal stress, and was enriched in Bacteroidota, Chloroflexota, filamentous Cyanobacteria, and Pseudomonadota associated with aerobic carbon fixation and oxidation, sulfur oxidation, and oxidative nutrient recycling.

### Community metabolic models reveal restructuring of nutrient exchange networks under geothermal stress

To quantify integrated community metabolism, we reconstructed four context-specific community metabolic models corresponding to the suite of complex geothermal stressors in 46 °C, 51 °C, 55 °C, and 61 °C biofilms, incorporating 87–130 MAGs and up to 4,514 reaction families. The models were constrained using transcriptionally informed reaction fluxes, species-level expression contributions, and measured physicochemical conditions.

Nutrient exchange analyses revealed strong stress-dependent restructuring of metabolic interactions among ecological guilds (Fig. 3A). At 46 °C, the co-consumption and cross-feeding of carbon-, nitrogen-, phosphorus- and sulfur-associated metabolites occurred predominantly between GIII and GII. GIII was the principal donor of carbon-, phosphorus-and sulfur-associated metabolites to GII. Although GI was less prominent in the exchange network, it served as the primary donor of nitrogen-associated metabolites to both GII and GIII. At intermediate temperatures (51–55 °C), nutrient exchange became increasingly multidirectional, while GII and GIII remained core to interactions. Across this GIII–GII-dominated interval, carbon-, nitrogen- and phosphorus-associated interactions shifted from co-consumption towards cross-feeding, whereas sulfur showed the opposite trend. At 55 °C, GI had emerged as the principal recipient of metabolites associated with all four elements. At 61 °C, GI became the dominant competitor for and donor of these metabolites, whereas GIII connectivity declined sharply.

**Figure 3.**
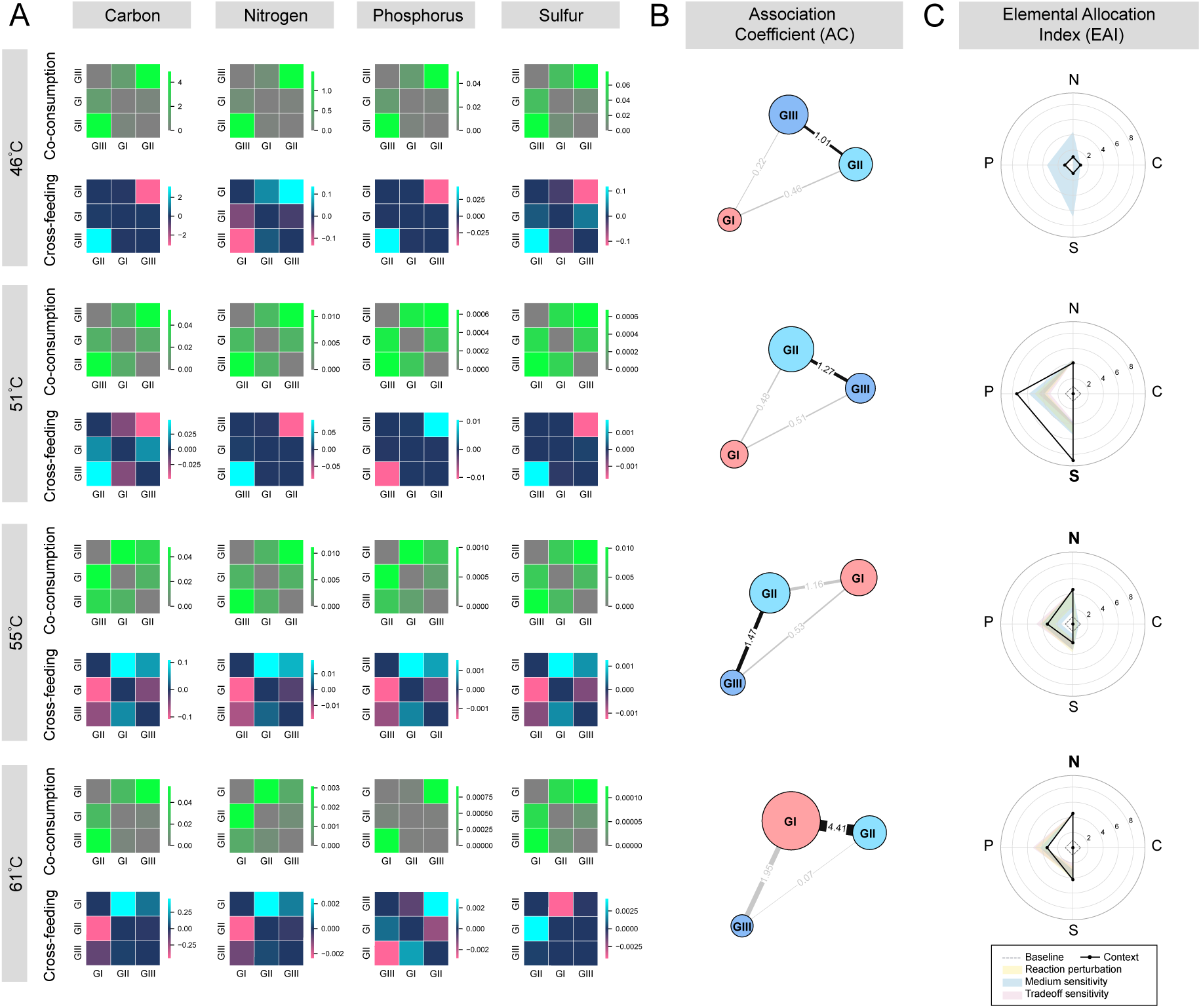
Metabolic interactions among ecological guilds across the hot spring geothermal gradient. **A)** Heatmaps showing co-consumption and cross-feeding interactions for carbon (C), nitrogen (N), phosphorus (P) and sulfur (S) fluxes among the three ecological guilds (GI–GIII) across thermal conditions (46 °C, 51 °C, 55 °C and 61 °C). Co-consumption matrices represent overlap in elemental resource utilization between guilds, whereas cross-feeding matrices represent predicted metabolic exchange and dependency relationships inferred from context-constrained community metabolic models. Colour intensity indicates the relative magnitude and direction of elemental flux interactions. Hierarchical clustering highlights temperature-dependent restructuring of metabolic connectivity among guilds. **B)** Association coefficient (AC) networks summarizing the overall strength of inter-guild metabolic interactions at each geothermal interval. Node size represents the relative contribution of each ecological guild to total community-wide metabolic activity, while line thickness and embedded numerical labels indicate the strength of inferred associations between guilds. **C)** Elemental allocation index (EAI) radar profiles showing relative modelled elemental demand, as shifts in external nutrient limitation pressure for C, N, P, and S in context-constrained CMMs relative to the unconstrained baseline CMMs. Bold type identifies the nutrient with the largest relative limitations. Shaded bands represent the interquartile ranges obtained from three sensitivity analyses (Supplementary Dataset 3), illustrating the robustness of inferred nutrient-allocation trends to uncertainty in transcriptomic constraints and parameter selection.

In recognition of the inherent challenges to quantitatively interpreting the balance between reciprocal cross-feeding and resource co-consumption, we calculated a novel metric, the Association Coefficient (AC). This expressed the balance of cross-feeding and co-consumption between taxon pairs, based on the geometric mean of bidirectional cross-feeding normalized by co-consumption intensity. The AC metric integrates reciprocal cross-feeding and competitive co-consumption into a single interaction metric, thereby reflecting the overall metabolic connectivity between ecological guilds. The AC metric was applied to our hot spring dataset (Fig. 3B), and revealed that at 46 °C, GIII formed the dominant interaction hub, and the strongest reciprocal interactions occurred between GII and GIII. Reciprocity increased further at 51 °C and 55 °C, as the GII–GIII interface remained the dominant reciprocal partnership across this intermediate thermal range. At 61 °C, the interaction peaked and shifted such that the strongest reciprocal occurred at the GI–GII interface, coinciding with dramatically reduced activity and connectivity of GIII. To assess model robustness, we conducted three sensitivity analyses targeting community–individual growth trade-off, environmentally detected nutrient-uptake bounds, and transcriptome-derived reaction constraints. Across logarithmically spaced AC values, the observed pattern remained consistent thus further supporting our conclusion (Supplementary Dataset 3).

Furthermore, to quantify the relative elemental allocation shifts for guilds across the geothermal gradient we calculated a second novel metric, the Elemental Allocation Index (EAI), by normalizing positive net elemental demand against the baseline unconstrained community simulation. The EAI thus quantifies how contextual environmental and transcriptomic constraints alter relative modelled elemental demand compared with the unconstrained baseline community model. Higher EAI values identify elements subject to greater relative context nutrient demand pressure. The EAI calculated for this system articulates the progressive shifts in community nutrient demand from sulfur- and phosphorous-associated limitation at lower temperatures, toward a pronounced switch to nitrogen-limitation at the highest temperature (Fig. 3C). Under the weakest geothermal stress in 46 °C samples, contextualization produced little overall change in elemental demand under the default parameterization. However, perturbing environmental nutrient-uptake bounds identified sulfur as the element most susceptible to depletion. An elevated sulfur demand gap became apparent at 51 °C, consistent with the distinct sulfur cross-feeding axis observed in the internal interaction network. Under higher geothermal stress at 55-61 °C, the system nutrient pressure transitioned from sulfur-dominated to nitrogen-dominated, indicating a major shift in metabolic allocation toward nitrogen-dependent processes under extreme geothermal stress. Sensitivity analyses varying the community–individual growth trade-off and transcriptome-derived reaction bounds retained nitrogen as the dominant element at high temperatures (Supplementary Dataset 3), while also identifying phosphorus as a potential secondary demand gap (Fig. 3C).

Together, these results suggest adaptation to geothermal stress in the hot spring biofilms involved coordinated restructuring of both internal nutrient interactions and external nutrient dependencies, with sulfur-associated interactions dominating under moderate geothermal stress and nitrogen limitation progressively emerging under elevated geothermal stress.

### Geothermal gradients redistribute elemental pathway flux among ecological guilds

Having established a geothermal stress-driven shift in nutrient limitation, we next investigated how these community-level constraints were realized through metabolic flux redistribution. We employed elemental pathway flux analyses to identify pronounced redistribution of biogeochemical function among guilds across the geothermal gradient (Fig. 4, Supplementary Dataset 3). At the whole-community level (indicated as “All”), carbon fixation at 46 °C was distributed across multiple pathways, with heterotrophic anaplerotic carbon fixation predominating and GII and GIII contributing most of the flux. At 55–61 °C, carbon metabolism was reorganized around a GI-dominated Calvin cycle–glycolysis–pyruvate oxidation axis, in contrast to the more distributed, TCA-associated carbon metabolism observed at lower temperatures. Nitrogen fixation was mediated largely by GII across the entire gradient, whereas nitrogen utilization displayed a distinct U-shaped response, with reduced amino acid utilization at intermediate temperatures approximately inversely tracking nitrogen fixation activity. Phosphorus utilization shifted from nucleotide-associated pathways at 46–51 °C to carbohydrate-associated pathways at 55–61 °C. Under greatest geothermal stress at 61 °C, GI dominated carbohydrate-associated processes involved in both phosphate release and metabolic incorporation, aligning phosphorus turnover with the GI-centred Calvin cycle–glycolysis–pyruvate oxidation axis. Sulfur acquisition was overwhelmingly derived from organic sulfur sources across the gradient; however, sulfur utilization underwent a major transition at 55-61 °C, where flux shifted away from amino acid biosynthesis toward sulfur reduction and sulfur oxidation bioenergetic pathways associated with GI, with sulfur reduction accounting for the larger share at 61°C. This redox shift was consistent with GI-associated Calvin-cycle flux, suggesting potential sulfur–carbon coupling under elevated geothermal stress.

**Figure 4.**
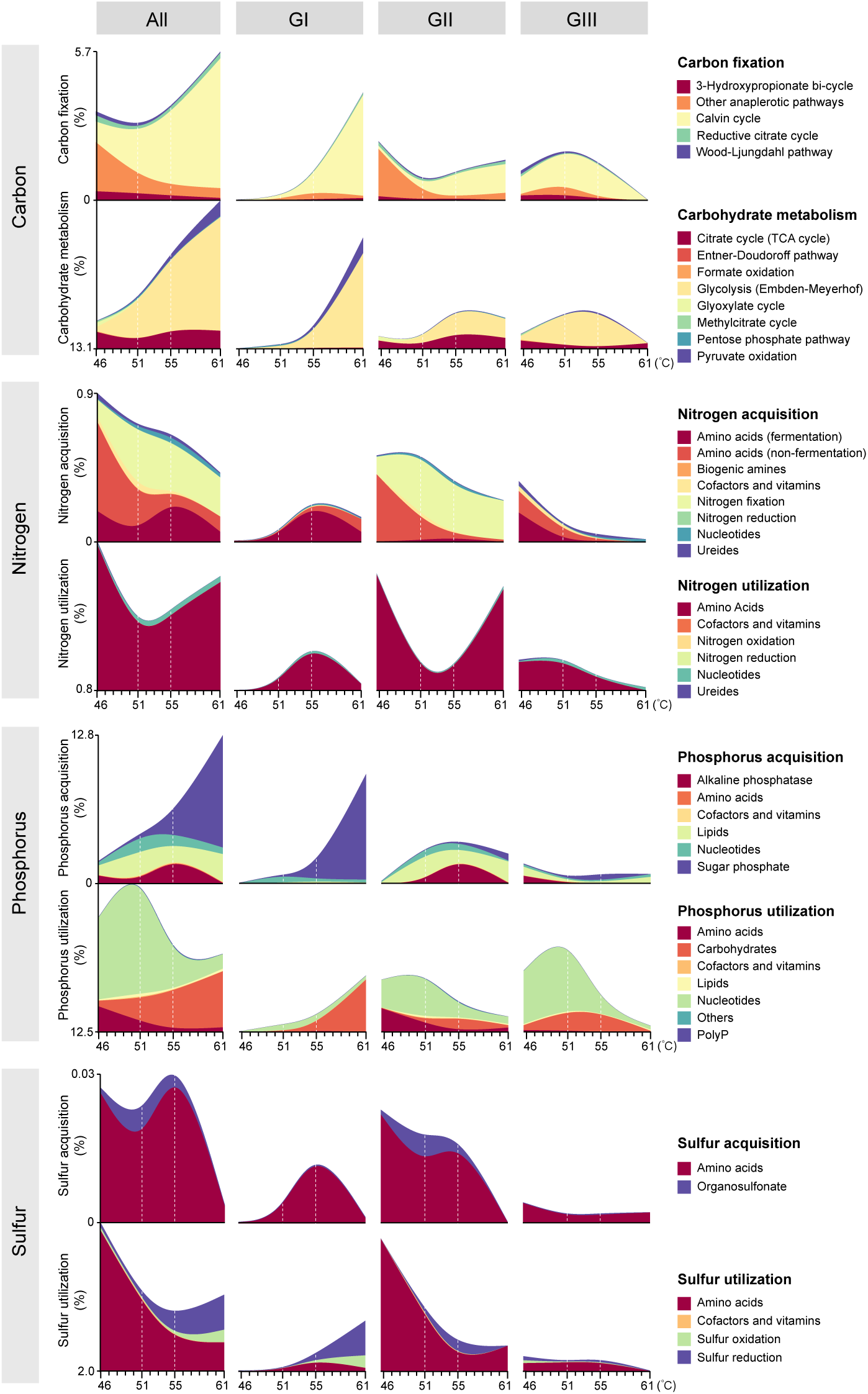
Restructuring of elemental pathway allocations in response to geothermal stress. Stress-dependent allocation of metabolic flux towards elemental acquisition and utilization pathways for carbon, nitrogen, phosphorus and sulfur cycling across the whole community (“All”) and the three ecological guilds (GI–GIII). Stacked area plots show the proportional contribution of individual pathways inferred from CMMs (as overall flux percentages) to elemental acquisition and utilization processes, including carbon fixation and carbohydrate metabolism, nitrogen acquisition and utilization, phosphorus acquisition and utilization, and sulfur acquisition and utilization. Functional allocation patterns shifted markedly with temperature and differed among guilds, indicating contrasting ecological strategies and resource partitioning under thermal stress.

### Geothermal stress reallocates electron flow toward catabolism and stress response

Next, we sought to identify community-scale electron allocation patterns, which we inferred from contextualized CMM flux solutions using manually curated functional electron-allocation modules. Reactions were classified into three major electron allocation modules corresponding to anabolism, catabolism, and stress response. This electron allocation model revealed a systems-level thermodynamic transition from anabolic biosynthesis toward catabolic energy conservation and stress-response metabolism with increasing geothermal stress (Fig. 5A, Supplementary Dataset 3). Within catabolism, flux from electron intermediates to organic compounds, sulfur oxidation, nitrate and nitrite increased, whereas flux to O₂ decreased. In contrast, electron flux to all anabolic destinations relatively declined, reaching its lowest level for organic synthesis, while stress-response allocation increased. Thus, communities under greater geothermal stress showed an electron-sink replacement from O₂-associated routes towards non-O₂ catabolic pathways, accompanied by reduced anabolic and increased stress-associated allocation.

**Figure 5.**
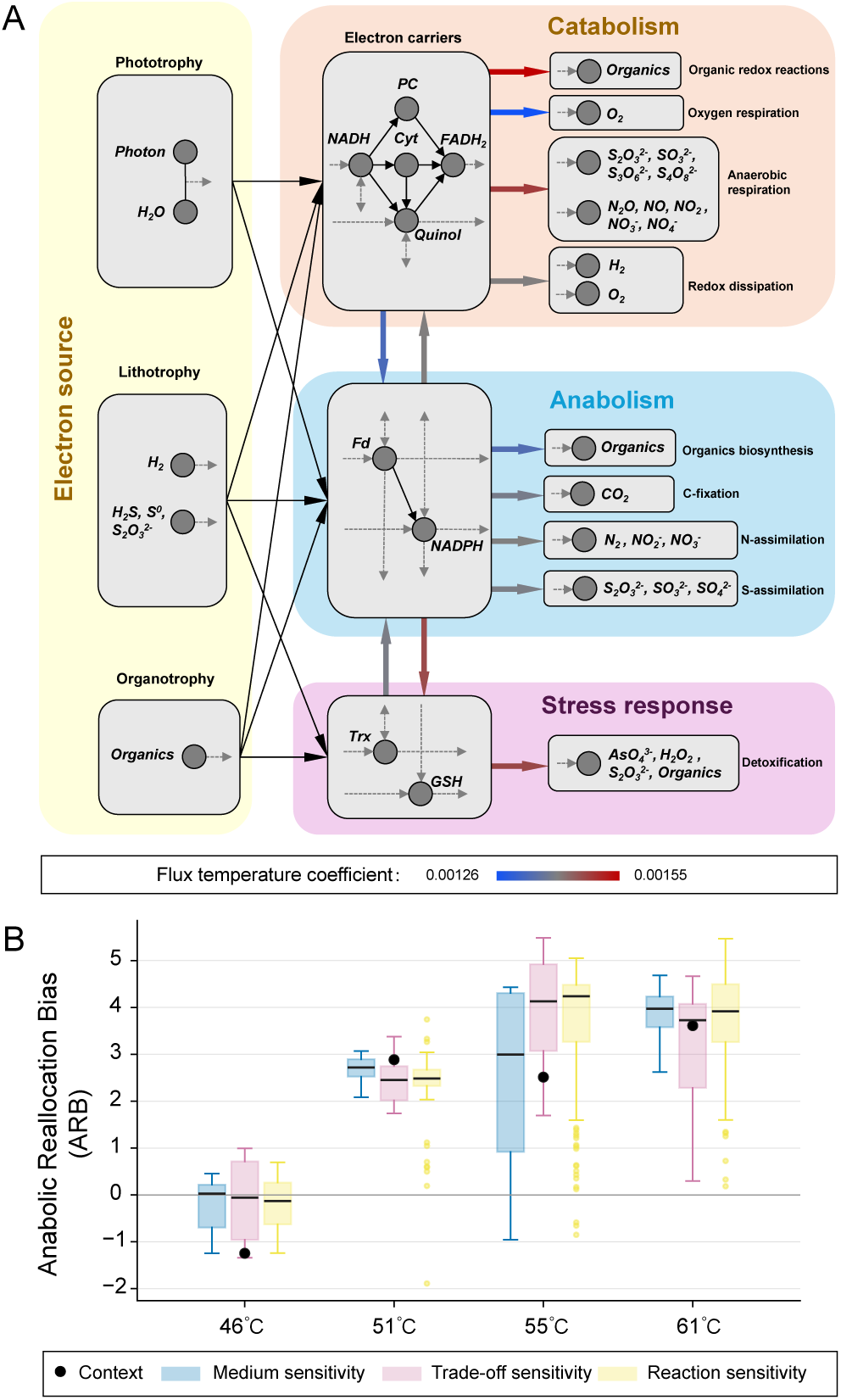
Electron allocation and metabolic energetic trade-offs in response to geothermal stress. **A)** Conceptual model of electron allocation across catabolic, anabolic and stress-response pathways inferred from context-constrained community metabolic models. The network diagrams depict electron flow from major energy inputs (oxygenic photosynthesis, organic oxidation, H_2_ oxidation, sulfur oxidation, and organics), through intermediate carriers (quinones, cytochromes, ferredoxins, NAD(P)H, thioredoxin, glutathione), to terminal sinks representing anabolic, catabolic, and stress-response pathways. Electrons derived from phototrophic, lithotrophic and organotrophic energy sources were distributed through intracellular electron carriers including NADH, quinols, cytochromes, ferredoxin (Fd), NADPH, thioredoxin (Trx) and glutathione (GSH), before ultimately being allocated to terminal anabolic, catabolic, and stress-response metabolic sinks. Red arrows indicate electron allocation pathways that increased with temperature, whereas blue arrows indicate pathways that decreased with temperature. Arrow colour intensity represents relative flux temperature coefficient. **B)** Anabolic reallocation bias (ARB) across the geothermal gradient (46–61 °C) for modelled (context) values predicted a progressive shift in electron allocation between anabolic, catabolic, and stress-associated metabolism (shown as black circles). At low stress (46°C) ARB was at or below zero indicating anabolic prioritization, whereas elevated geothermal stress resulted in a pronounced switch to positive ARB values indicating re-prioritization of metabolism towards catabolic and stress response metabolism. Values derived from the model were robust under three separate sensitivity analyses (coloured boxplots, Supplementary dataset 3).

In order to summarize this transition in a single quantitative measure, we introduce the Anabolic Reallocation Bias (ARB) metric to summarize this transition. Here, reactions within the three electron allocation modules (anabolism, catabolism, stress response) were additionally grouped into directional transitions linking function, such that positive ARB values indicate net electron reallocation away from anabolic investment toward catabolic or stress-associated functions, whereas negative ARB values indicate preferential reallocation toward anabolic metabolism. For our dataset, the ARB shifted from slightly negative values at 46 °C toward strongly positive values at 51–61 °C, a pattern retained across all three sensitivity analyses (Fig. 5B, Supplementary Dataset 3), indicating progressive reallocation of electron flux away from anabolic investment and toward catabolic and stress-associated functions (Fig. 5B, Supplementary Dataset 3). This transition was consistent with increasing prioritization of maintenance-oriented catabolism and stress-response pathways ahead of growth-oriented biosynthesis. Together, these results support our observations for the response in nutrient exchange and elemental flux, and suggest a coordinated guild-level reorganization of overall nutrient exchange, elemental flux, and electron allocation across the geothermal gradient.

## Discussion

### Geothermal gradients restructure microbial ecosystems through guild-level metabolic partitioning

Our integrated metatranscriptome-guided metagenome-resolved systems biology approach revealed that geothermal gradients restructure hot spring biofilms through coordinated reorganization of ecological guilds, metabolic interactions, and bioenergetic allocation. Rather than observing a simple taxonomic turnover along the geothermal gradient, we identified emergent guild-level metabolic strategies that partitioned ecosystem function according to thermal and redox conditions. These guilds transcended phylogenetic boundaries and instead reflected convergent ecological strategies shaped by energy availability, nutrient exchange, and electron acceptor utilization. The low-stress GIII guild was characterized by oxidative heterotrophic recycling and diverse carbon fixation pathways, whereas the high-stress GI guild became increasingly dominated by reductive sulfur-linked metabolism, thermophilic carbon fixation, and stress-adapted energy conservation. Between these extremes, GII functioned as a transitional nutrient-processing guild that maintained intermediary nutrient transfer and metabolic coupling across the geothermal gradient. These findings demonstrate how geothermal stress reorganizes microbial ecosystems through coordinated functional partitioning among co-occurring guilds rather than through replacement by isolated specialist taxa alone^34^.

### Metabolic interaction networks reveal increasing specialization under geothermal stress

The interaction networks and AC metric further revealed that increasing temperature progressively altered the organization of metabolic dependency within the ecosystem^15,29^. The community under lower geothermal stress displayed relatively distributed metabolic interactions centred on the oxidative GIII guild, consistent with broad nutrient recycling and comparatively high functional redundancy^34^. In contrast, intermediate geothermal stress resulted in maximal reciprocity and ecological integration, suggesting that moderate geothermal stress enhances metabolite exchange and cooperative nutrient transfer among guilds. Under the highest geothermal stress conditions, however, the network became increasingly centralized around the GI thermophilic guild, while overall reciprocity and connectivity declined. These observations suggest that elevated geothermal stress constrains ecosystem complexity and reduces metabolic redundancy^35^, leading to tighter coupling of specialized metabolic functions. Such restructuring is consistent with ecological theory predicting that environmental filtering increasingly favours specialized and energetically efficient metabolic strategies under environmental stress^36^, and offers a mechanistic explanation for the observed speciation of niche specialists in hot springs^37^.

The shift toward cooperation under extreme conditions aligns with the stress-gradient hypothesis in ecology^38^, which posits that facilitative interactions increase as stress intensifies. Our findings are supported by co-occurrence observations in other hot spring systems^15^, and our results offer a mechanistic explanation for such observations by providing quantitative, flux-based validation. Our MICOM-based approach was implemented with specific focus on demonstrating species-level contribution scaling^39^, allowing guild interactions to be interpreted within specific environmental contexts. The framework supports multi-omics integration and can be further refined as richer constraint data become available. Benchmarking against SMETANA^40^ showed broad agreement in interaction architecture: Even without condition-specific environmental, transcriptomic or species-scaling constraints, SMETANA recovered a GI-centred interaction potential that most closely resembled the core pattern observed under the highest geothermal stress (Supplementary Dataset 3). Overall, we propose that these features make the approach described here a practical proxy for capturing how microbial functional interactions are reorganized under environmental stress.

### Sulfur-linked metabolism and electron reallocation dominate ecosystem function under elevated geothermal stress

A major emergent property of the geothermal transition was the increasing dominance of sulfur-linked metabolism and associated electron management pathways. At lower stress levels, carbon fixation and carbohydrate metabolism were distributed across multiple pathways including anaplerotic carbon fixation, the Calvin cycle, the 3-hydroxypropionate bicycle and TCA-associated metabolism. However, this distributed configuration gave way to a GI-dominated Calvin cycle–glycolysis–pyruvate oxidation axis increasing geothermal stress. Phosphorus metabolism shifted in parallel from nucleotide-associated utilization at to carbohydrate-associated flux. By contrast, nitrogen represented the dominant elemental demand gap in the thermal ends. The intensified nitrogen demand gap may constrain the conversion of carbon and phosphorus turnover into anabolic products and biomass accumulation which was mirrored by electron allocation modelling and our articulation of the ARB metric, showing that under high geothermal stress, electron allocation shifted from anabolism towards non-O₂ catabolism and stress maintenance, including sulfur-linked energy metabolism.

Complementing this C-N-P reorganization, sulfur metabolism became increasingly important across the coupled thermal–redox transition. Sulfur compounds provide versatile electron donors and acceptors capable of supporting both oxidative and reductive energy metabolism under fluctuating redox conditions^41^, and sulfur cycling has long been recognized as a defining feature of thermophilic ecosystems ^42^. In our study, sulfur utilization underwent a pronounced transition at the highest geothermal stress, from amino acid-associated biosynthesis toward sulfur oxidation and sulfur respiration coinciding with the emergence of GI as the dominant metabolic guild. Coupled with increased electron allocation to respiratory and stress-response pathways, these findings suggest that sulfur metabolism increasingly functions as a central energetic axis supporting redox balancing and maintenance metabolism under geothermal stress^43^. Importantly, this transition occurred concurrently with reduced metabolic diversity and increasing centralization of ecological interactions, indicating that this hot spring ecosystem may operate under increasingly constrained energetic regimes near the upper thermal limits for a given guild. Laboratory studies have shown that carbon fixation is more thermolabile than photosystem activity in hot spring cyanobacteria^44^, and so our study provides an explanation for how alternate routes for reducing power may arise for photosynthetic biofilms under geothermal stress.

### Energetic constraints underlie ecological thresholds in hot spring communities

The observed increase in anabolic reallocation bias (ARB) toward catabolism and stress response further supports the hypothesis that geothermal stress imposes substantial energetic and thermodynamic constraints on microbial metabolism^45,46^ and communities^47,48^. Although thermophilic taxa are adapted to high temperatures, our findings suggest that communities above ∼55–60 °C at Sembawang Hot Spring increasingly prioritize maintenance and redox stabilization above growth-oriented biosynthesis. This provides a potential mechanistic explanation for the sharp geothermally-defined ecological transitions commonly observed in hot springs, e.g., ≤ 60 °C for most filamentous cyanobacteria^49^ and ≤ 75 °C for photosynthetic bacteria^7,50^, where small increases in geothermal stress can drive abrupt shifts in community composition and function. Reduced metabolic redundancy and increasing reliance on tightly coupled reciprocal interactions may additionally reduce ecosystem resilience under extreme conditions, potentially rendering communities under elevated geothermal stress more sensitive to environmental perturbation.

The modelled shifts define a two-phase metabolic strategy along the geothermal gradient where with increasing stress, biomass investment declines as electrons are redirected toward energy maintenance, alternative respiration, and detoxification. This represents a possible ecological tipping point from growth- to survival-oriented metabolism, a transition hypothesized in other hot spring systems^15,51^, but here substantiated with quantitative estimates from community-scale flux modeling based upon multi-omics integration. The shifts observed under geothermal stress corresponded with a successive decline of predicted secondary and primary productivity, which is consistent with ecological observations indicating that stressed communities undergo progressive nutrient limitation and functional collapse^52^. At the community level this may emerge as a generalizable feature of environmental stress response in microbial systems, analogous to that proposed for oligotrophic marine systems where metabolic constraints are imposed by nutrient stress^53^.

### Systems biology reveals emergent principles of microbial ecosystem organization

More broadly, this study demonstrates the value of integrating genome-resolved metagenomics, transcriptomics, environmental chemistry, and community metabolic modelling to understand emergent ecosystem function in extreme environments. Traditional taxonomic or functional gene distribution approaches alone cannot readily resolve how microbial communities partition metabolism, exchange nutrients, and reorganize energy flow under environmental stress^3^. By combining multi-omics data with systems-level modelling, we were able to identify functional ecological guilds, quantify nutrient exchange networks via the Association Coefficient (AC), infer shifts in nutrient allocation via the Elemental Allocation Index (EAI), and quantify electron allocation under stress via the Anabolic Reallocation Bias (ARB) metric across the environmental gradient. We propose that these novel metrics help to define the overall drivers of community turnover and advance our understanding of community-level adaptation beyond the occurrence of niche specialist and generalist taxa inferred from taxonomic studies in hot springs^37^. The findings suggest that microbial community metabolism is best understood as an emergent network property arising from coordinated interactions among ecological guilds rather than as the sum of independent organismal functions ^29^. This community level insight on adaptation to geothermal stress complements a recent study where highly plastic metabolism in laboratory cultures of individual hot spring bacteria indicated species level adaptive traits^41^. Overall, this is congruent with the hypothesis that metabolic plasticity and functional redundancy in communities may be co-selected in response to environmental stress^36^.

Despite the interpretive power of our framework, it is not without limitations. Transcription-guided flux constraints approximate metabolic capacity but cannot fully capture post-transcriptional regulation, enzyme kinetics, or rapid allosteric responses, thus potentially underestimating short-term metabolic plasticity near steep physicochemical gradients. In addition, the models assume relatively stable substrate supply at the biofilm–water interface, whereas microscale stratification^54^, stochastic disturbance^55^, diel^56^ and seasonal^55^ variability also likely influence electron donor availability and nutrient transfer *in situ*. Future work integrating time-resolved metatranscriptomic and proteomic constraints, together with direct experimental measurements of metabolite uptake and redox dynamics, may improve estimates of microbial activity and flux partitioning, and enable further validation of the metabolic modelling framework.

Overall, we have demonstrated how an integrated systems biology approach can reveal novel insight on the putative ecological response of microbial biofilms to geothermal stress. It will be interesting to extend this in future studies to broader extreme environmental gradients in hot springs, as well as for other microbial communities facing environmentally driven shifts in metabolic organization, for example in biofilms exposed to climate-related stress or in engineered bioprocesses. We anticipate that the novel Association Coefficient (AC) to quantify cross-feeding and co-consumption, the Elemental Allocation Index (EAI) to quantify shifts in nutrient allocation, and the Anabolic Reallocation Bias (ARB) to quantify electron reallocation between metabolic strategies, may become useful tools for articulating such responses in diverse ecosystems.

## Methods

### Environmental samples

Photosynthetic biofilms were sampled along a 46°C–61°C geothermal gradient at the circumneutral Sembawang Hot Springs, Singapore (N 1.434460, E 103.822000). Biofilms were collected from each of four temperature-defined intervals (46, 51, 55, 61 °C; N = 12), and the biomass was transferred into screw-cap tubes containing 500 µL RNAlater preservative (ThermoFisher), stored on ice in darkness during transport, and kept at −20°C until processed. Abiotic water variables were measured as follows: temperature (digital thermocouple, Fluke); pH, ORP, electrical conductivity, and fluoride (ion-selective electrode, Hach); hydrogen sulfide (H_2_S; colorimetric assay, Hach); total organic carbon (TOC) (TOC analyzer, Shimadzu); ammonium, nitrate, nitrite, sulfate (Dionex ion analyzer, ThermoFisher); phosphate (colorimetric test, AMS); bromide and chloride (ion chromatography, Metrohm); and metal ions, including cadmium, calcium, iron, lithium, magnesium, sodium, and potassium (ICP-OES, Agilent). Temperature and H_2_S were measured on-site, and all other variables were analyzed in the laboratory immediately after sampling.

### DNA and RNA sequencing

DNA was extracted from biofilm samples using the PowerLyzer PowerSoil Kit (Qiagen). For RNA extraction, samples were ground in liquid nitrogen using a sterilized mortar and pestle, processed using the TRIzol RNA isolation kit (ThermoFisher), and depleted of rRNA using the Ribo-Zero Plus kit (Illumina). Sequencing library preparation was performed using PE150 kits and sequencing carried out using the NovaSeq 6000 platform (Illumina). DNA and RNA quality was examined using a Bioanalyzer (Agilent). Quality control, including adapter removal and trimming, was performed using Fastp (v0.23.1)^57^ with default settings. Human sequences were removed by mapping to the hg38 reference genome using Bowtie2 (v2.4.5)^58^. For metatranscriptomic data, rRNA sequences were filtered using SortMeRNA (v4.3.2)^2^ using its default database. After processing, an average of 13.4 ± 1.0 GB per metagenomic sample and 4.9 ± 1.7 GB per metatranscriptomic sample were retained

### Reconstruction, annotation, and expression-based quantification of MAGs

Clean paired-end metagenomic reads were merged using BBMerge (v.36.2)^38^ and assembled into contigs using metaSPAdes (v3.15.4)^59^. Binning was performed using MetaBAT (v0.24.1)^60^ and MaxBin (v2.2.7)^61^, yielding 476 bins, which were dereplicated at ≥95% ANI using DAS Tools^62^. MAG quality was assessed using CheckM (v1.1.9)^63^, and those with <50% completeness or >10% contamination were excluded^64^, resulting in 140 medium-to high-quality MAGs for downstream analysis. Taxonomic classification and phylogeny were determined using GTDB-Tk (release 2.4.0)^65^, and phylogenomic trees were visualized in iTOL (v7.2.2)^66^. Open reading frames were predicted using Prokka (v1.14.5)^67^. Functional annotation was performed using eggNOG-mapper (v2.1.9)^68^ and Rapid Annotation using Subsystem Technology (RAST)^69^. The relative abundance of MAGs in samples was estimated using CoverM (v0.6.1)^70^. The meta-data for all MAGs is reported in Supplementary Dataset 1.

Metatranscriptomic activity of MAGs was quantified using the a genome-wide gene expression (GE) dataset with average expression of 92 conserved prokaryotic housekeeping genes identified with UBCG (v3.0)^71^ (Supplementary Dataset 1), and used as a multi-omic qualification input for downstream analysis and modeling because it better reflects strain-specific activity and provides a more accurate representation of the community level metabolic state^72^ than DNA abundance based scaling. Housekeeping genes were selected because metatranscriptomic datasets typically exhibit highly uneven transcriptional coverage across MAGs, with expression concentrated in active loci and limited coverage across non-expressed genomic regions^73^. In addition, lineage-specific accessory genes can bias whole-genome expression estimates^74^. RNA abundance profiles were analyzed using the MIntO (v2.0.0) pipeline^75^ and normalized as transcripts per million (TPM).

### Identification of expression-defined ecological guilds

Expression-defined ecological guilds were inferred from MAG-level GE profiles across 12 sample-level observations. Pairwise Spearman correlations were calculated for all MAG pairs (N = 140; 9,730 pairs), generating a complete edge table containing Spearman’s Rho, raw P values, and Benjamini–Hochberg adjusted Q values. Because nominal Spearman P values were not used as the sole basis for guild definition, the primary co-expression network was defined as a weighted positive-correlation network retaining MAG–MAG edges with Rho > 0, with edge weights set to Rho. Louvain, Walktrap, and multilevel modularity detections were then applied to this weighted network to define the primary guild framework using igraph (v1.0.0)^76^. Partition robustness was evaluated using silhouette analysis and adjusted Rand index (ARI) comparisons, and the most concordant modularity partitions were used as the primary framework for defining ecological guilds (Supplementary Dataset 2). The primary partition identified three major ecological guilds (G_I_-G_III_), with MAG097 and MAG075) did not affiliate with any of the ecological guilds. These MAGs were nonetheless included in downstream community-scale modeling to avoid exclusion bias.

To assess robustness to network construction, modularity detection was repeated across alternative edge definitions, including positive edges with raw P ≤ 0.05, positive edges with BH-adjusted Q ≤ 0.05, strong-correlation thresholds (Rho ≥ 0.5, 0.6, and 0.7), top-positive edge subsets, leave-one-temperature-group-out stable edges, and within-temperature bootstrap-stable edges. Leave-one-temperature-out stable edges were required to remain positive in all recalculations and to have a median leave-one-out Rho ≥ 0.5. Bootstrap-stable edges were defined as positive edges retained in at least 80% of 200 within-temperature bootstrap iterations, with strong-correlation support also required in at least 80% of iterations. Alternative guild assignments were compared with the primary Louvain partition using adjusted rand index (ARI), normalized mutual information (NMI), and assignment concordance after optimal label matching (Supplementary Dataset 2).

### Construction of genome-scale metabolic models (GEMs)

Draft genome-scale metabolic models (GEMs) were constructed for each MAG using CarveMe (v1.4.1)^77^ based on universal models for Bacteria and Archaea. Because these models incompletely capture sulfur metabolism, despite the abundance of sulfur compounds in hot spring environments, we manually incorporated sulfur-related reactions^78^ into selected MAGs using COBRApy (v0.29.0)^79^ (Supplementary Dataset 3). Genes associated with these reactions were identified by clustering MAGs with reviewed protein sequences from UniProt Release 2023_02 via OrthoFinder (v2.5.4)^80^, and reactions were added when corresponding proteins were detected. Automated gap-filling was then applied to ensure that the GEM model reflected each strain’s biological traits^81^. For instance, photosynthetic bacteria were required to grow using light as the sole energy source and inorganic carbon as the sole carbon source; chemolithoautotrophs using inorganic substrates (e.g., hydrogen, reduced sulfur, or iron) as sole energy sources^82^ and inorganic carbon as the sole carbon source; nitrogen-fixers using nitrogen as the sole nitrogen source; and anaerobes without oxygen. The basal nutrient requirements for constraining growth models of MAGs in a photoautotrophy-driven biofilm community were adapted from a universal bacterial basal medium^81^, supplemented with light and thiosulfate^83^, arsenite, and arsenate^84^ which are compounds common in circumneutral, moderately sulfidic hot springs. To meet specific gap-filling needs modifications were made to light, oxygen, carbon, and nitrogen sources, and simulating both aerobic and anaerobic conditions. Strains not meeting these criteria for growth were simulated in a more flexible rich medium consisting of the hot spring basal medium plus 172 metabolites predicted as exchangeable among draft GEMs based on SMETANA (v1.1.0)^40^ in the global mode. The resulting photoautotrophic basal plus cross-feeding media under aerobic and anaerobic conditions represented potential environmental resources or cross-fed metabolites. All downstream analyses, therefore, assume that the curated GEMs adequately represent the metabolic capacities of their corresponding MAGs; and we acknowledge that residual uncertainty in lineage-specific reaction coverage may affect estimates of modelling outputs. Detailed curation statistics, gap-filling summaries, and MEMOTE quality scores for all curated GEMs are provided in Supplementary Dataset 3.

### Construction and simulation of metatranscriptome-based community metabolic models (CMMs)

The community metabolic models (CMMs) were constructed from GEMs using MICOM (v0.36.3)^39^ with the CPLEX (v22.1.1) solver in Python (v3.10.11). Four CMMs were developed, one for each environmental sample set. Community members with detectable GE values and normalized abundances above the MICOM tolerance threshold were incorporated into the CMM topology, with GE used directly as the community scaling parameter. After filtering, we assembled 128 (at 46 °C), 126 (at 51 °C), 130 (at 55 °C), and 87 (at 61 °C) GEMs with MEMOTE score 80 ± 4% (mean ± s.d.) into CMMs for the biofilms at each temperature. The initial CMMs were unconstrained, with reaction flux bounds left at default values and with GE used as the community scaling parameter. Protein-associated reaction constraints were derived from RNA abundance using RIPTiDe (v3.4.81)^85^. Constraints associated with MAGs showing near-zero omics signals with discordant DNA–RNA detection were removed, thereby minimizing noise from weak or inconsistent multi-omics evidence. Protein-associated reaction constraints of these eligible MAGs subsequently incorporated into the CMMs as transcriptome-informed reaction bounds (Supplementary Dataset 3), producing context-specific models constrained by the metatranscriptomic data.

Community growth simulations were performed to evaluate both baseline and context-specific growth of CMMs using the cooperative tradeoff framework (tradeoff = 1). Baseline growth was simulated in an unconstrained medium with unrestricted metabolite availability. Hot spring-specific growth was simulated using a metatranscriptome-constrained model and a medium representative of the Sembawang hot spring (Supplementary Dataset 3), based on the basal medium used in auto-gap-filling and supplemented with site-specific water chemistry. Medium parameterization was constrained by measured metabolites: molecular nitrogen was set as the sole nitrogen source due to the absence of detectable dissolved inorganic nitrogen and phosphate, although below detection limits, was permitted minimal uptake flux using the micom.media.minimal_medium algorithm to satisfy basal growth requirements. For redox-sensitive metabolites, oxygen and iron fluxes were determined by hydrogen sulfide measurements: anoxic conditions (O_2_ = 0) with exclusive Fe^2+^ influx under sulfidic conditions versus oxic conditions with unrestricted Fe^2+^/Fe^3+^ influx when H_2_S was absent. Cofactors and vitamins, essential for CCM functionality, were included in the modeling medium during automatic gap-filling, but their influx was restricted to 1 ×10⁻⁶ of the default allowable rate (0.001 mmol gDW^−1^ total biomass h^−1^) to minimize their impact. All other detected compounds were assumed to be in excess with unrestricted uptake bounds, and undetected compounds were excluded. Unless otherwise specified, downstream ecological metrics were calculated from community simulation fluxes after GE-weighted aggregation across MAG-specific reactions.

### Elemental interaction analysis

Potential metabolic interactions among community members were inferred from interspecies exchange reactions in the contextualized CMM flux solutions. Pairwise elemental interaction fluxes for carbon (C), nitrogen (N), phosphorus (P), and sulfur (S) were inferred using the micom.interaction framework. Interaction fluxes were subsequently aggregated at metabolic groups level based on ecological guild assignments.

To quantify the balance between reciprocal cross-feeding and resource co-consumption, we calculated an Association Coefficient (AC) between taxon pairs based on the geometric mean of bidirectional cross-feeding normalized by co-consumption intensity:

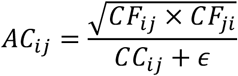

where *CF_i_*_j_ represents directional cross-feeding flux from taxon *i* to taxon *j*, *CC_i_*_j_ represents co-consumption flux between taxa *i* and *j*, and *ε* = 10^−7^ was included to avoid division-by-zero artifacts throughout all downstream ecological metric calculations unless otherwise specified. Sensitivity analysis across *ε* varying from 10^−6^to 10^−9^ confirmed the robustness of metric rankings, with all absolute deviations remaining under 1% (Supplementary Dataset 3). Higher AC values indicate stronger reciprocal metabolite exchange relative to shared co-consumption, whereas lower AC values indicate dominance of resource competition over bidirectional metabolic exchange. As an orthogonal benchmark, guild-level metabolic interaction potentials and resource overlap were additionally evaluated using SMETANA (Supplementary Dataset 3). Because SMETANA does not natively incorporate transcriptome-weighted activity scaling, these analyses were performed using guild-level interaction summaries without transcriptome-informed activity weighting.

### Elemental limitation analysis

Community-level elemental demand for C, N, P, and S was quantified as the net elemental uptake flux derived from medium-associated exchange reactions in the contextualized CMM simulations. To quantify relative modelled elemental allocation shifts across environmental conditions, we calculated an Elemental Allocation Index (EAI) by normalizing positive net elemental demand against the baseline unconstrained community simulation:

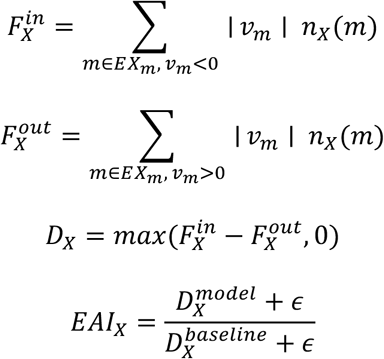

where *v_m_* is the exchange flux of metabolite *m*, *n_X_*(*m*) is the number of atoms of element *X* in metabolite *m*, and *D_X_* represents the positive net environmental demand for element *X*. Thus, EAI quantifies how environmental and transcriptomic constraints alter relative elemental demand compared with the unconstrained baseline community model. Values of *EAI_X_* > 1 indicate increased net relative demand for element *X* relative to the baseline simulation, whereas *EAI_X_* < 1 indicates reduced net demand. Relative elemental allocation profiles across C, N, P, and S were additionally visualized in Python using the matplotlib (v3.9.2) package with polar coordinate projection.

### Electron allocation analysis

Community-scale electron allocation patterns were inferred from contextualized CMM flux solutions by assigning electron-carrier-associated reactions a priori to functional modules based on their established biochemical roles. These modules comprised three major electron allocation modules: anabolism (ANA), catabolism (CATA), and stress response (STRESS). To characterize electron reallocation among physiological modules, reactions were additionally grouped into directional transitions linking anabolism with catabolism and stress-response functions (Supplementary Dataset 3), from which the Anabolic Reallocation Bias (ARB) was calculated as:

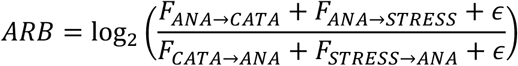

where *F* represents the aggregated directional reallocation flux associated with each directional transition. Positive ARB values indicate net electron reallocation away from anabolic investment toward catabolic or stress-associated functions, whereas negative ARB values indicate preferential reallocation toward anabolic metabolism.

### Sensitivity analyses

To evaluate the robustness of inferred elemental interaction and electron-allocation patterns, we performed three sets of sensitivity analyses each conditional on the four context-specific CMM. First, we varied the MICOM cooperative-tradeoff parameter across its full range from 0 to 1 at increments of 0.5, thereby testing the sensitivity of model predictions to different balances between community-level optimality and individual member growth requirements. Second, to evaluate the influence of unconstrained default medium values, we applied a default medium-bound gradient to key default-bounded medium components representing major elemental substrates (CO₂, bicarbonate, sulfate, nitrate, nitrite, ammonium, and N₂), energy sources (light, hydrogen, and reduced sulfur), and redox status (O₂), varying upper uptake bounds from 5 to 1000 across 12 levels. Third, to evaluate the influence of uncertainty in internal reaction constraints, we perturbed protein-associated reaction constraints in contextualized CMMs within their feasible reaction-flux intervals.

We conducted reaction-level sensitivity analysis by perturbing transcriptome-informed constraints on protein-associated reactions within their feasible flux intervals. Perturbations were conducted at the reaction-family level to preserve shared functional constraints across MAG instances carrying the same metabolic reaction. For each realization, 5% of eligible reaction families were randomly selected and perturbed using Latin hypercube sampling (LHS; sample size = 100), introducing sparse transcriptome-informed variability into reaction constraints while preserving community feasibility and the overall metabolic structure of the contextualized CMMs. Downstream ecological metrics, including elemental interaction profiles and electron allocation patterns, were recalculated for each sensitivity analysis (Supplementary Dataset 3). Robustness statistics, including medians, confidence intervals, and interquartile ranges were subsequently calculated across all sensitivity ensemble types to evaluate the stability of ecological interaction and metabolic allocation patterns under transcriptome-informed constraint parameterization uncertainty.

## Data availability

Sequence data for all metagenomes, metatranscriptomes, and metagenome-assembled genomes are publicly available in NCBI under BioProject PRJNA1106483. The reaction parameters for metabolic models are archived at Zenodo DOI: 10.5281/zenodo.20132606. Supplementary data (Supplementary Datasets 1-3) in support of the findings are available online. All codes specific to this study are archived at github: https://github.com/luodanli-research/MetaContext-CMM-pipeline.

## Acknowledgments

We thank the Singapore National Parks Board for facilitating hot spring sampling under permit NP/RP21-126-1. We also thank Dolyce Hong Wen Low (Duke-NUS Medical School) for advice and assistance with RNA extractions.

## Author contribution statement

D.L., P.K.H.L., and S.B.P. designed the study. C.G. and S.B.P. performed field and laboratory experiments. D.L., C.G., P.K.H.L., and S.B.P. performed bioinformatic analysis. D.L. developed, implemented, and validated the metabolic models with input from P.K.H.L. and S.B.P. Interpretation of the findings and manuscript preparation were undertaken by D.L., P.K.H.L., and S.B.P.

## Ethics declaration

The authors declare no competing interests.

## Supplementary Datasets

**Supplementary Dataset 1.** Quality metrics, mapping rates, taxonomy abundance, and functional annotation of 140 metagenome-assembled genomes.

**a)** Quality metrics, taxonomy, DNA abundance, and genome-wide gene expression (GE) of 140 metagenome-assembled genomes (MAGs).
**b)** Proportion of quality-controlled metagenomic reads mapped to the 140 reconstructed MAGs, indicating representativeness across samples.
**c)** Number of genes categorized into RAST functions in each MAG.

**Supplementary Dataset 2.** Ecological groups and adaptive functions related to aqueous geochemistry gradients.

**a)** Aqueous geochemistry of the hot spring geothermal gradient.
**b)** Robustness of ecological guild detection across modularity algorithms.
**c)** Robustness of ecological guild assignments to alternative co-expression network definitions.

**Supplementary Dataset 3.** Gap-filling procedures for genome-scale models, including manual reactions and media for automated gap-filling, and parameterization of community-scale metabolic models using aqueous geochemistry-based media and metatranscriptome-derived reaction constraints.

**a)** Gap-filling methods and quality check of curated 140 genome-scale models (GEMs).
**b)** Parameterization of community-scale metabolic model (CMM) media by aqueous geochemistry.
**c)** Reaction constraints applied to CMMs across environmental gradients.
**d)** Sensitivity analysis statistics for nutrient interaction (AC and EAI) and electron allocation (ARB) metrics.
**e)** SMETANA benchmark of internal interactions between ecological guilds.
**f)** Reaction sets used for elemental pathway allocation and anabolic reallocation bias (ARB) analyses.

